# Physics-constrained neural ordinary differential equation models to discover and predict microbial community dynamics

**DOI:** 10.1101/2025.07.08.663743

**Authors:** Jaron Thompson, Bryce M. Connors, Victor M. Zavala, Ophelia S. Venturelli

## Abstract

Microbial communities play essential roles in shaping ecosystem functions and predictive modeling frameworks are crucial for understanding, controlling, and harnessing their properties. Competition and cross-feeding of metabolites drives microbiome dynamics and functions. Existing mechanistic models that capture metabolite-mediated interactions in microbial communities have limited flexibility due to rigid assumptions. While machine learning models provide flexibility, they require large datasets, are challenging to interpret, and can over-fit to experimental noise. To overcome these limitations, we develop a physics-constrained machine learning model, which we call the Neural Species Mediator (NSM), that combines a mechanistic model of metabolite dynamics with a machine learning component. The NSM is more accurate than mechanistic or machine learning components on experimental datasets and provides insights into direct biological interactions. In summary, embedding a neural network into a mechanistic model of microbial community dynamics improves prediction performance and interpretability compared to its constituent mechanistic or machine learning components.

**Significance statement:** Microbial communities drive essential biological processes in every ecosystem. Predicting microbial community dynamics and functions and uncovering their interaction networks are challenging due to their complexity and emergent interactions. We develop a framework that combines biological knowledge with machine learning to accurately predict community and metabolite dynamics. This approach outperforms existing ecological and machine learning models by capturing environmentally mediated interactions while providing interpretable insights into microbe-metabolite interactions. Our framework enables the rational design and control of microbial communities for applications in medicine, agriculture and biotechnology

## Introduction

Microbial communities play critical roles in nearly every ecosystem, performing essential functions in systems ranging from human health [49] to waste valorization [46]. Within an ecosystem, microbial community growth dynamics are largely governed by resource availability as well as time-dependent production and consumption of metabolites [2]. Although interactions between microorganisms change over time, in presence of other species, and across different environmental contexts, existing modeling approaches are often based on simplified mechanisms that do not capture this context dependency [2]. Conversely, machine learning modeling of microbial communities provides flexibility, but is challenging to interpret and can require large amounts of training data to accurately learn system behaviors [3, 51]. Developing flexible, yet interpretable, models that can accurately predict the temporal behaviors of microbial communities based on their initial composition and environmental inputs is essential to understand and design microbial communities.

The generalized Lotka-Volterra (gLV) model is a widely used dynamic model and assumes that species dynamics are governed by pairwise interactions with all constituent community members [52, 41, 9]. Therefore, the gLV has limited flexibility for capturing higher-order interactions and does not capture metabolite dynamics. Microbe-effector models are a broad class of mechanistic models that represent interactions between species and metabolites and are typically based on either mass-action or Monod kinetics [39]. Within this class of models, the MacArthur consumer resource (CR) assumes mass-action kinetics where species growth rates are proportional to the product of their abundance and resource concentration [2]. A key limitation of microbe-effector models is the assumption of constant interaction coefficients, which limits their ability to capture complex interactions between species and metabolites or state and time dependent shifts in metabolite production and consumption. For example, the consumption and release rates of metabolites in microbial communities can change as a function of time, leading to complex dynamical behaviors [44, 19]. Finally, since the full metabolite dynamics shaping microbial communities are not known and/or challenging to measure experimentally, microbe-effector models are typically not used for data-driven applications. Machine learning models offer a powerful approach to capture the complexity of microbial community dynamics, often surpassing the prediction performance of traditional mechanistic models on experimental data. For example, dynamic machine learning models, such as recurrent neural networks (RNNs), have been used to predict changes in species abundances and metabolite concentrations over time [3, 51]. However, as discrete-time models, RNNs are constrained to making predictions at fixed time intervals, making them difficult to apply to irregularly sampled data and limiting their ability to interpolate between time points. Neural ordinary differential equations (NODEs) address this limitation by modeling dynamics in continuous time [11]. However, existing implementations of NODEs applied to ecological systems have either focused on prediction at a single time point or have relied on highly time-resolved data to predict dynamics. For instance, NODEs have been used to predict metabolomic profiles and microbiome compositions at a fixed time point [53, 37]. By contrast, NODEs have been applied to capture both simulated and experimental dynamic ecological data that included a single initial condition, with 10-100 time series measurements that can provide rich information for estimating the direction of change [1, 8, 7]. However, experimental data from synthetic microbial communities often consist of potentially hundreds of distinct initial conditions, each with only a few (*<*5) time series measurements of species abundances or metabolite concentrations [17, 52, 12, 3, 20]. In sum, the prediction performance of NODEs on data from synthetic microbial community experiments presents challenges that have not yet been evaluated.

Although machine learning models such as RNNs and NODEs have the potential to capture complex dynamics, the lack of a mechanistic structure necessitates learning the foundational physical principles of species growth and metabolite production entirely from limited experimental data. An additional limitation of machine learning models compared to mechanistic models is that they can be difficult to interpret and require post hoc methods such as Shapley Additive exPlanations (SHAP) [32] or Local Interpretable Model-agnostic Interpretations (LIME) [43]. While these approaches can provide useful insight into how a model translates experimental variables such as species initial abundance to predictions of biological functions, they require additional computation after training a model and cannot reveal direct interactions [3, 13]. Combining physics-based mechanistic models and machine learning models can result in improved flexibility over entirely mechanistic models and prediction performance over machine learning models [26]. Machine learning models can be physically constrained by incorporation of a learning bias that penalizing instances where the machine learning predictions violate physical laws during training [24]. To ensure that model predictions are physically relevant, incorporation of inductive bias involves embedding a physics-based structure into the machine learning model. A key advantage of the incorporation of inductive bias is that it does not require data to learn the physical laws of the system. Further, embedding mechanistic models into machine learning models can preserve interpretability of the mechanistic component. There are many existing frameworks that combine mechanistic and machine learning models. Universal ordinary differential equations [40] (UODE) add neural network outputs to mechanistic ordinary differential equation models to learn unknown dynamics. However, because the neural network is added rather than integrated into the mechanistic model, the structure of UODEs does not prevent physically unrealistic predictions. Recently, modifications to enforce non-negativity for applications in biochemical systems have been proposed [38]. A separate framework that is becoming widely used to model physical systems is physics-informed neural networks (PINNs) [24]. PINNs use a neural network to learn a function whose partial derivatives satisfy a known partial differential equation (PDE). While this approach can provide an efficient way to approximate solutions of PDEs, this method is generally useful in cases where a mechanistic model is known in advance, such as fluid mechanics [10]. Methods such as UODEs, which seek to learn gaps in existing mechanistic models from experimental data are especially promising for modeling microbial communities, where mechanistic knowledge of system behavior is more limited.

To leverage the strengths of mechanistic and machine learning models of synthetic microbial communities, we embed a mass-action kinetic model of metabolite production and consumption as a physical scaffold into a neural ordinary differential equation model. The mechanistic component forces the model to obey physical properties such as species-mediated metabolite production and consumption while the machine learning component captures potentially non-linear dependence of metabolite production and consumption rates on community and environmental context. This physics-integrated machine learning model (Neural Species Metabolite or NSM) significantly outperforms both entirely mechanistic models and entirely machine learning models, especially when the amount of training data is limited. Further, we incorporate a scalable Bayesian inference algorithm for estimating model parameters and quantifying uncertainty [6]. By analyzing uncertainty in parameter estimates, we show that confidently identified parameters corresponding to the mechanistic component of the model recovers known interactions between species and metabolites using simulated and experimental data. In sum, we show that carefully integrating mechanistic and machine learning models for microbial communities can substantially enhance predictive performance, enforce physical constraints, and yield interpretable model parameters.

## Results

### A physics-informed machine learning model for metabolite and microbial community dynamics

Mechanistic ODE models have the benefit of a model structure that enforces physical relevance. However, the complexity of microbial communities makes it challenging to define a mechanistic model that fully captures the complex system dynamics. By contrast, while more flexible machine learning models can adapt to complex experimental data, they can also be more challenging to train on limited and noisy data without the risk of over-fitting. We propose to combine the strengths and overcome limitations of mechanistic and machine learning models by fusing a neural network with a mechanistic model of metabolite production and consumption (Fig. 1). The mechanistic component enforces metabolite consumption rates that are proportional to species abundance and production rates that are proportional to species growth rate. This structure is specifically tailored to model primary metabolites, which are produced while species are actively growing [48, 2]. Incorporating this mechanistic structure captures a dominant mode of interaction between species involving competition for limited resources and cross-feeding of metabolites [15]. To capture a broader range of interaction mechanisms such as toxin production, the neural network takes time-dependent species abundance, metabolite concentrations, and static experimental inputs (e.g., initial pH) to predict the rate of change of species and metabolites. With the combined flexibility of the neural network and the physical groundwork of the mechanistic model, the hybrid model can adjust the rate of change of metabolites and species without having to learn the foundational principle that metabolites are produced and consumed by species.

**Figure 1:**
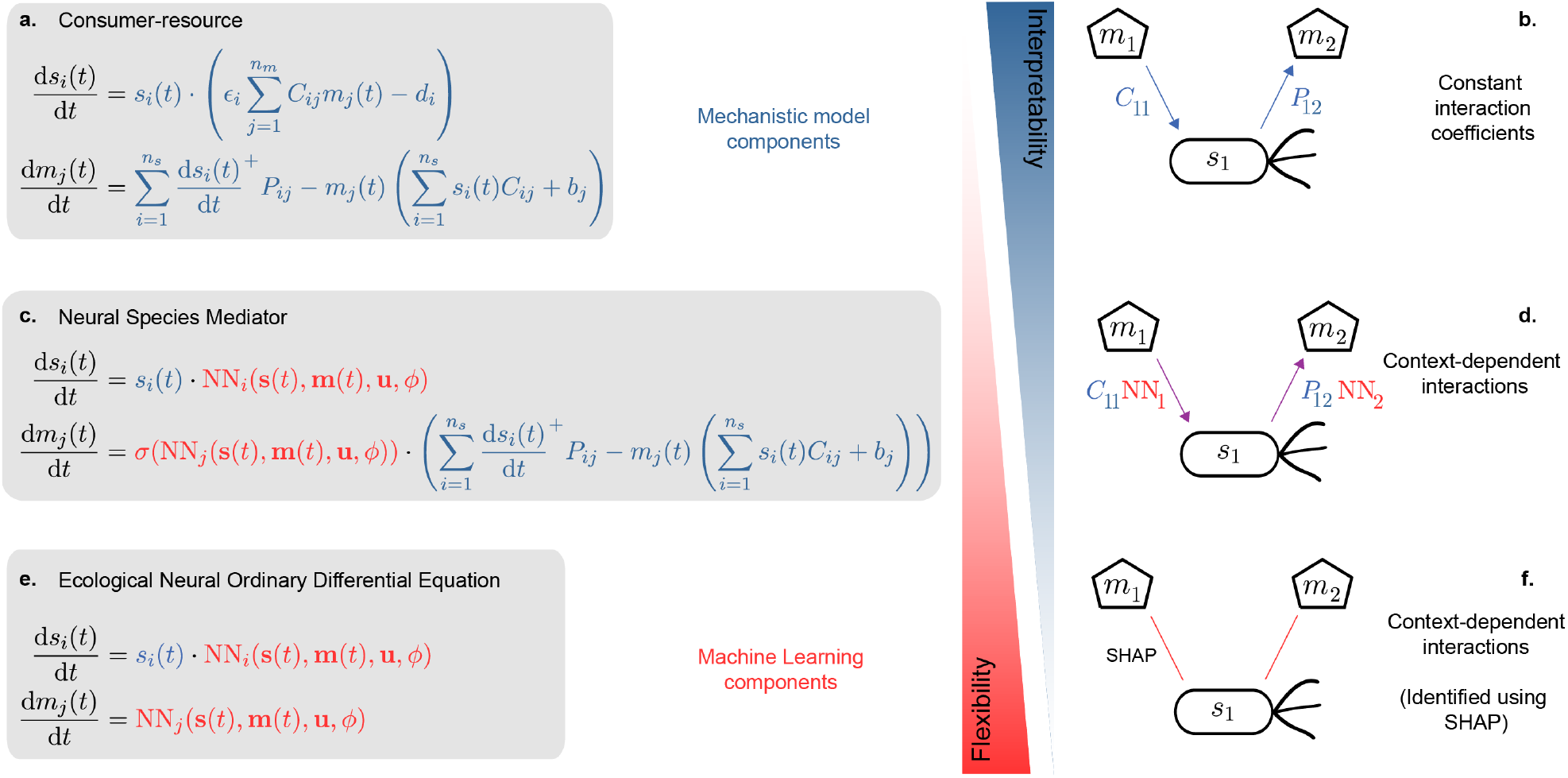
Neural species mediator (NSM) model. (a.) The consumer-resource model is a standard ecological model of species growth dynamics governed by competition and cross-feeding of metabolites. (b.) The model parameters of the consumer-resource model quantify the rate that species consume and produce metabolites. While the assumption of fixed consumption and production coefficients enables straightforward interpretation, the model cannot capture context-dependent species-metabolite interactions. (c.) The NSM uses a neural network to predict species growth rates and to modify the consumer-resource model of metabolite dynamics. (d.) The neural network takes as input species abundance and metabolite concentration to predict how consumption and production rates should be modified, enabling context dependent species-metabolite interactions while preserving the interpretable consumption and production rate parameters. (e.) The eNODE relies on a neural network to predict species and metabolite dynamics. Unlike a standard NODE, the eNODE incorporates the multiplication of species abundance, which forces the model to predict growth rates of zero for species whose abundance is zero. (f.) The eNODE does not have parameters that are directly interpretable and requires post-hoc methods such as SHAP to identify species-metabolite interactions.

We denote the time-dependent vector of species abundances as 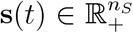, metabolites as 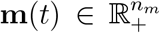, and additional system inputs as 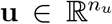. We consider the case where system inputs **u** are static, although the framework could be extended to incorporate time-dependent inputs. We refer to the proposed model as the neural species mediator (NSM), which is given by the following set of coupled differential equations,

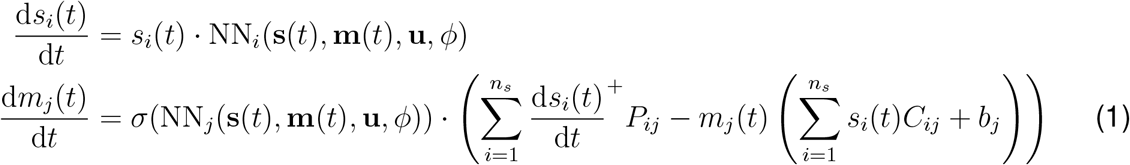

where *P*_*ij*_ ∈ ℝ_+_ is the strictly-positive growth-associated rate that species *i* produces metabolite *j, C*_*ij*_ ∈ ℝ_+_ is the strictly-positive rate that species *i* consumes metabolite *j*, and *b*_*j*_ ∈ ℝ_+_ is the constant degradation rate of metabolite *j*. We use rectified linear unit function, *a*^+^ = max(0, *a*), to model mediator production as proportional to species growth rate (i.e. mediators are assumed to not be produced during stationary or death phase) [48, 33]. The structure of the mechanistic component can easily be modified to account for biomass associated production where mediators are produced during stationary and death phase (Eq. 4). A feed-forward neural network with parameters *ϕ* (Methods) has an output dimension of *n*_*s*_ + *n*_*m*_. For *i* = 1, …, *n*_*s*_, the neural network output, NN_*i*_(**s**(*t*), **m**(*t*), **u**, *ϕ*), captures the growth rate of species *i*. A key challenge in integrating mechanistic and machine learning components is ensuring that the machine learning model complements rather than overrides the mechanistic understanding in making predictions. For *j* = *n*_*s*_ + 1, …, *n*_*s*_ + *n*_*m*_, the product of the neural network output *σ*(NN_*j*_(**s**(*t*), **m**(*t*), **u**, *ϕ*)) and the mechanistic model of metabolite production predicts the rate of change of metabolite *j*. We use the softplus activation function, *σ*(*a*) = log_*e*_ (1 + *e*^*a*^), to force the output of the neural network to be positive. Unlike direct exponentiation, the softplus function grows linearly with large input values, which helps prevent numerical overflow and improves stability during training. Because the neural network output is strictly positive, the sign (positive or negative) of the rate of change of metabolites is determined entirely by the mechanistic component. Consequently, the NSM cannot rely exclusively on the neural network to predict metabolite dynamics. Forcing the neural network output to be positive also prevents metabolite concentrations from becoming negative since the consumption rate of metabolites decreases to zero as metabolites approach zero. By contrast, these properties do not necessarily hold when the neural network is added to the mechanistic model, as proposed by UODEs (Eq. 22) which can produce physically unrealistic negative predictions [1, 40]. In sum, the NSM integrates a neural network to provide flexibility while preserving the structure of the mechanistic model, ensuring predictions adhere to physical principles. Capturing the mechanistic logic where metabolites are either produced or not produced at all depending on whether the metabolite producing species is present or absent is not straightforward for neural networks, which generally produce smooth variation in outputs in response to variation in inputs [5].

We refer to the species component of the NSM as an ecological neural ordinary differential equation (eNODE) model. The key modification is the multiplication of species abundances into the neural network in the right-hand side of the ODE. This simple constraint forces species with zero-valued initial conditions to remain zero and prevents non-zero species trajectories from becoming negative. Incorporating this physical constraint greatly improves the model’s ability to fit experimental data. Similarly, the addition of the mechanistic component of the NSM prevents predictions of metabolites from becoming negative, which provides an analogous advantage for the prediction of metabolites. To demonstrate the benefit of incorporating these constraints into the model architecture, we evaluated the fit of the NODE, eNODE, UODE, and NSM to simulated data from a ground truth consumer-resource model (Fig. S1). The ground truth consumer-resource model includes eight species and six metabolites, where each species can consume or produce metabolites. Simulated data was generated by randomly sampling 32 combinations of species and metabolites initial conditions, sampling the dynamics at four time points over a 32-hour period, and then introducing Gaussian noise to simulate variability in measurements (Methods). Models that are easier to train show a faster drop in prediction error (root-mean-square-error or RMSE) on the training set. Additionally, models with increasing depth and width of the neural network can achieve a lower fitting error. As the number of hidden units (network width) and layers (network depth) increases, the NODE and eNODE match the fit achieved by the UODE and NSM but require more training/learning iterations. By contrast, the UODE and NSM were able to fit the data in fewer training iterations, demonstrating that combining mechanistic and machine learning models can make it easier to train and enables the use of a simpler neural network component. Further, the NSM and UODE display a longer computational time per epoch than eNODE and NODE. Further, the computation time generally increases as training proceeds (Fig. S2). This trend is consistent with the numerical integration solver, which may require more computation time as the solver adjusts time step sizes to maintain integration accuracy [47]. In sum, incorporating mechanistic components into a NODE results in faster and more stable training without sacrificing the ability to fit the data.

### The NSM outperforms existing models in prediction tasks of experimental data

#### The eNODE outperforms the gLV in prediction of species abundances

The generalized Lotka-Volterra (gLV) model is a widely used model of microbial community dynamics that captures pairwise interactions between all constituent community members [50, 52]. However, this model cannot capture higher-order interactions between species. Machine learning methods such as the long short-term memory RNN (LSTM) have shown improved prediction performance and computational scalability compared to the gLV [3]. The restriction of RNNs to discrete-time intervals makes it challenging to train on irregularly sampled time-series data and impossible to make predictions at intermediate time intervals. However, NODE models have not yet been benchmarked for their ability to capture the dynamics and functions of synthetic microbial communities compared to widely used dynamic models. Therefore, we evaluated the prediction performance of the eNODE and gLV on an experimental data set composed of combinations of 11 bacterial species, including four different *Clostridioides difficile* (CD) strains (Fig. 2a) [50]. Samples that included co-cultures were characterized at 12 and 24 hours, and monoculture samples were measured at 3-hour intervals from 0 to 24 hours. Species absolute abundance was determined by multiplying relative abundance based on 16S rRNA gene sequencing by the total community biomass based on absorbance at 600 nm [12]. Using 10-fold cross-validation, the eNODE displayed significantly greater prediction performance for 5 species, *Bacteroides thetaiotaomicron* (BT), *Clostridium hiranonis* (CH), *Bacteroides uniformis* BU, *Desulfovibrio piger* (DP), and the MS008 CD strain, according to the correlated coefficients test. For the remaining species, the prediction performance of the NSM was not significantly different than the gLV (Fig. 2). Both models used the same train and test partitions of the data and the variational inference training algorithm (Methods). For the eNODE, each training fold was further partitioned into 10 folds and cross-validation on this nested set of folds was used to identify the neural network architecture that maximized prediction performance. Possible architectures included either one or two layers, each with either 5, 10, or 15 nodes in each layer. The improvement in prediction performance of the eNODE compared to the gLV is an important demonstration that more flexible approaches are required to capture the complexity of microbial community dynamics.

**Figure 2:**
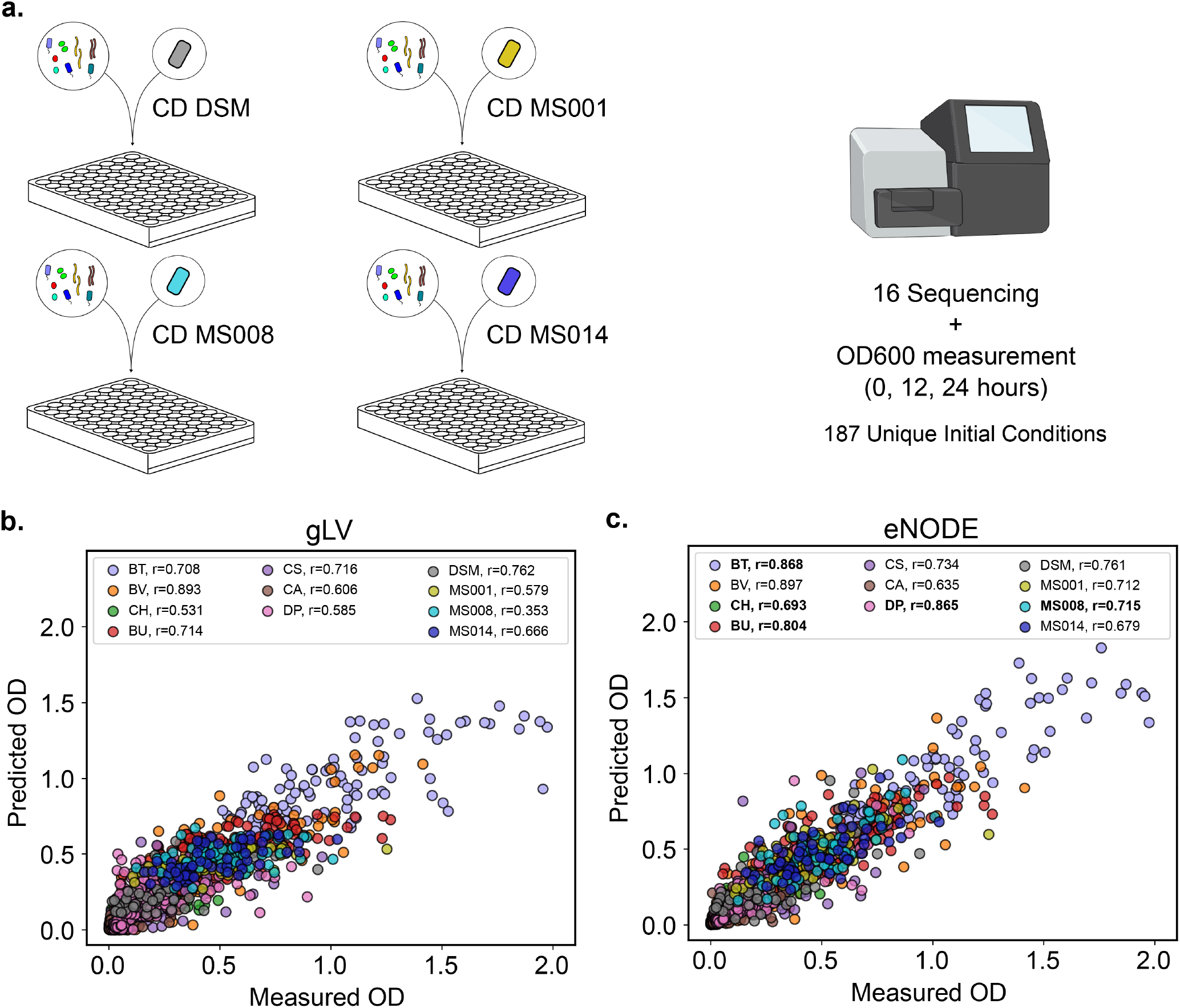
NSM/eNODE outperforms gLV in prediction of species. a. Schematic of experimental design. Briefly, four genetically distinct *Clostridium difficile* (CD) strains were individually inoculated into communities with varying numbers of diverse human gut bacteria and species abundances were measured at 12 and 24 hours after inoculation. The data set included 187 unique initial conditions. b. Generalized Lotka-Volterra (gLV) model predictions of species abundance based on 10-fold cross-validation. Prediction performance was evaluated using the Pearson correlation coefficient (r) between measured and predicted species abundances. c. eNODE model predictions of species abundance from the same 10-fold cross-validation splits as in panel b. Correlation coefficients that were significantly greater according to the correlated coefficients test for the eNODE compared to the gLV are bold.

#### The NSM outperforms the CR and eNODE models in prediction of metabolites

To evaluate the NSM’s ability to predict metabolites, we performed an experiment to investigate how interactions between two bacterial species, *Faecalibacterium prausnitzii* (FP) and *Desulfovibrio piger* (DP), influence the production of a metabolite, hydrogen sulfide, over the course of 37 hours. Previous studies have shown that DP reduces sulfate to produce hydrogen sulfide, which can contribute to inflammatory bowel disease in high concentrations [34]. Previous work has also demonstrated that DP inhibits butyrate production [12]. To study the dependence of species growth and metabolite production on initial substrate concentrations, the experiment data included 10 monoculture (FP or DP) and 10 community (FP and DP) conditions under varying media formulations where the concentrations of glucose, lactate, and sulfate were varied in each condition. To evaluate the prediction performance of the NSM and its mechanistic and machine learning components, we used a leave-one-out cross-validation approach on held-out samples in the FP-DP data set (Fig. 3a). The CR, NSM, UODE, and eNODE models were each trained using the same training algorithm using the same training data. We used a nested cross-validation approach where the full data set was partitioned into 10 folds. Each subset was further partitioned into 9 folds and cross-validation on this set was used to identify the neural network architecture that maximized prediction performance. Possible architectures included either one or two layers, each with either 5, 10, or 15 nodes in each layer. The CR and NSM predictions of hydrogen sulfide are forced to be non-negative due to their model structure. However, the eNODE and UODE models are capable of predicting negative hydrogen sulfide concentrations (Fig. 3c). The CR model was significantly worse than the NSM and eNODE at prediction of FP, DP, and hydrogen sulfide (Fig. 3d). While species predictions were not significantly different between the eNODE, UODE, and NSM, predictions of hydrogen sulfide from both the UODE and NSM were significantly more correlated with measured values compared to eNODE (p *<* .001, correlated coefficients test). In sum, models that combined mechanistic and machine learning components (UODE and NSM) outperformed a purely mechanistic model (CR) and a machine learning model (eNODE) at prediction of hydrogen sulfide.

**Figure 3:**
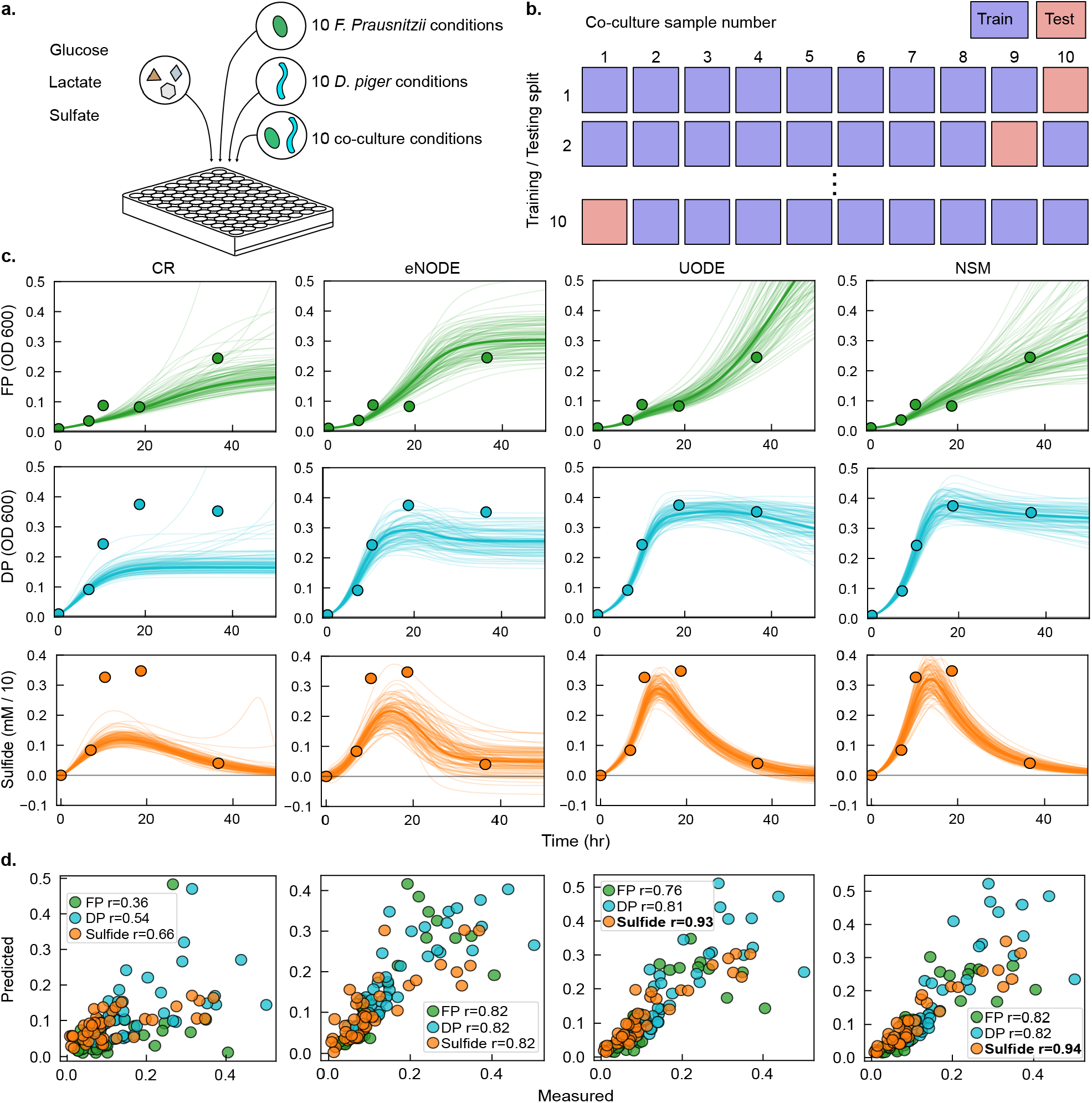
NSM and UODE outperform CR and eNODE in prediction of hydrogen sulfide. a. Species abundance (OD 600) and hydrogen sulfide concentration (mM) from monoculture and co-cultures of *F. Prausnitzii* (FP) and *D. Piger* (DP), each with varying initial concentrations of glucose, lactate, and sulfate, were measured at 7, 10, 19, and 37 hours after inoculation. b. A schematic to illustrate leave-one-out cross-validation, where each co-culture sample was subjected to held-out testing. c. CR, eNODE, UODE, and NSM predictions of a held-out condition are shown using 100 samples drawn from the posterior predictive distribution with the solid line representing the average model prediction. Scatter plot points represent measured values. d. Prediction performance of FP, DP, and hydrogen sulfide by the CR, eNODE, UODE or NSM demonstrates that the UODE and NSM significantly outperform the eNODE model in prediction of sulfide (p *<* .001, correlated coefficients test).

#### The NSM requires less data to make accurate predictions of metabolites

The incorporation of a physical constraint in the NSM could reduce the amount of training data required to make accurate predictions by enforcing generalizable rules such as species mediated production or consumption of metabolites. To test this possibility, we used a combined data set from Clark et al. 2021 [12] and Baranwal et al. 2022 [3] that comprised 757 combinations of 25 bacterial species where species abundances and 4 metabolites (acetate, butyrate, lactate, and succinate) were measured at either 48 hr or 16, 32, and 48 hr (95 samples). We evaluated the prediction performance of the eNODE, UODE, and NSM model when trained on a fraction of the data and then tasked the model to predict all remaining samples. To account for variation due to random sampling in the data, we repeated the process of randomly generating training and testing sets 50 times. To compare prediction performance between each model, we used the correlated coefficients test (Methods) to compare the average Pearson correlation over the 50 trials. For consistency in comparison between models and training data sizes, all models used a neural network with a single hidden layer with 20 nodes as this architecture was sufficiently flexible for accurate predictions of metabolites. The NSM consistently outperformed the eNODE over the entire range of training percents for all metabolites and displayed a significantly higher prediction performance than UODE and eNODE when the amount of training data was limited (Fig. 4). This result demonstrates that incorporating a physical constraint can improve generalizability of the model, especially when training data are limited. When the amount of training data is not limited, the physical constraint becomes less beneficial for prediction performance as the eNODE is able to learn the system from data.

**Figure 4:**
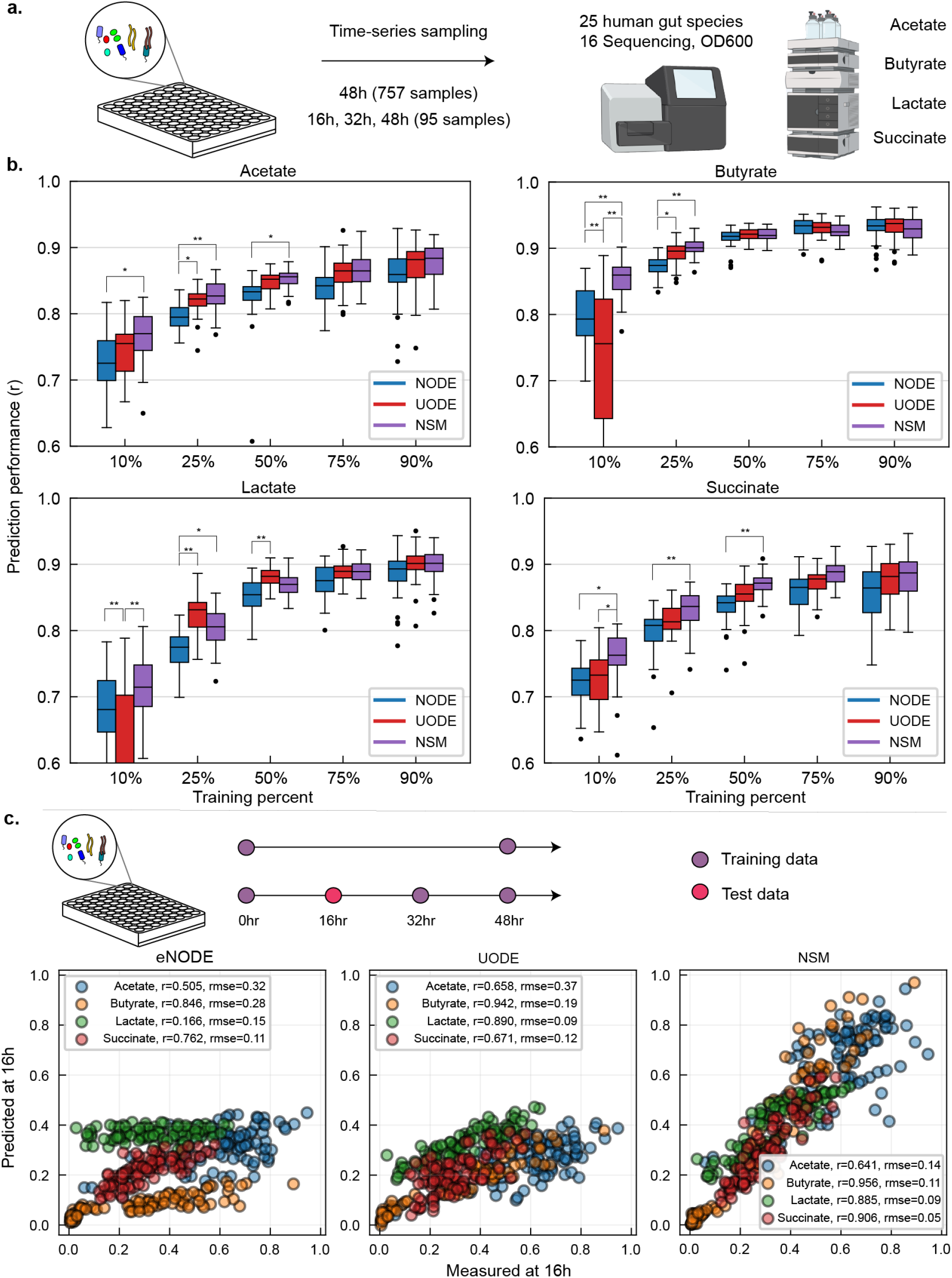
NSM is more accurate than UODE and eNODE given less training data. a. Experimental data of species abundances and concentrations of acetate, butyrate, lactate, and succinate from co-culturing combinations of 25 health-relevant human gut bacteria was used to train the NSM. Estimated consumption and production parameters provide insights into the rate that each species consumes or produces metabolites. b. Prediction performance (Pearson correlation coefficient) of metabolites using the eNODE, UODE, and NSM was compared as the amount of training data was increased from 10% to 90% of the full data set. The average Pearson correlation taken over 50 randomly generated training and testing partitions was compared using the correlated coefficients test (* p*<* .01, ** p*<* .001). c. Comparison of the eNODE, UODE, and NSM ability to predict measured metabolite values at 16 hours after training on all data excluding the 16-hour time point.

Despite comparable prediction performance between the eNODE, UODE, and NSM models given sufficient training data (Fig. 4b), model predictions of the eNODE and UODE showed more erratic trajectories in-between measurements of metabolites compared to the NSM when both models were trained on the entire data set (Fig. S3). This implies that the structure of the NSM promotes physical relevance when interpolating. To further investigate the predictions in between time points, we partitioned the data into training and test sets where measurements of species and metabolites at the 16-hour time point were withheld from training and reserved for the test set. When tasked with predicting metabolite values at the 16-hour time point, both the eNODE and UODE frequently underestimated measured values, yielding higher RMSE (root-mean-square-error) compared to the NSM (Fig. 4c). This result demonstrates that incorporating physical constraints can improve predictions of conditions beyond the training set even when provided with a high amount of training data.

#### The NSM identifies direct interactions between species and metabolites

A major limitation of machine learning models is that they can challenging to understand how the model predicts experimental dependent variables given independent variables (i.e. interpretability). To address this limitation, explainable artificial intelligence methods have been proposed to explain how machine learning models make predictions. One of the most widely used approaches is called Shapley additive explanations (SHAP) [31], which assigns an additive contribution of each model feature (independent variable) to the model prediction (dependent variable). The sum of the contributions of each feature equals the predicted value, and the magnitude of the contribution reflects the feature’s importance. It is important to note that SHAP contributions depend on the experimental condition being analyzed. Therefore, feature importances determined by SHAP do not reflect a global property of the system. Therefore, SHAP values are computed and analyzed over multiple conditions and provide insights into context-dependent interactions. Finally, a limitation of SHAP contributions is that they do not represent direct causal relationships between independent and dependent variables.

Incorporating a mechanistic component into machine learning models can provide insights into the biological system since the parameters have physical meaning. For example, the parameter *P*_*ij*_ represents the growth-associated rate that species *i* produces metabolite *j* (Fig. 1). The mechanistic model parameters are fixed values after training and therefore do not depend on the experimental condition. Unlike SHAP values, the coefficients in mechanistic models can capture direct and causal relationships.

To validate the ability of the NSM to identify direct interactions between species and metabolites, we fit the NSM to simulated data from a ground truth consumer-resource model with known metabolite consumption and production rates (Fig. 5a). The ground truth system represents combinations of eight species and six metabolites, where each species produces and consumes different subsets of metabolites resulting in a system governed by competition and cross-feeding. To determine whether the neural network component of the NSM would disrupt recovery of ground truth consumption and production rates, we fit the NSM to simulated species and metabolite data from the ground truth model (Fig. 5b). Simulated data was corrupted with zero-mean Gaussian noise to mimic variation in experimental measurements (Methods). As the number of training conditions (defined by the initial condition) increased from 16, 32, and 64, the ability of the NSM to accurately recover consumption and production rates approached a perfect correlation (Fig. 5c). Compared to the UODE, the NSM was able to recover parameters with greater accuracy, especially when the amount of training data was limited (Fig. 5c). Therefore, incorporation of the neural network in the NSM does not disrupt the ability to estimate physically relevant mechanistic model parameters.

**Figure 5:**
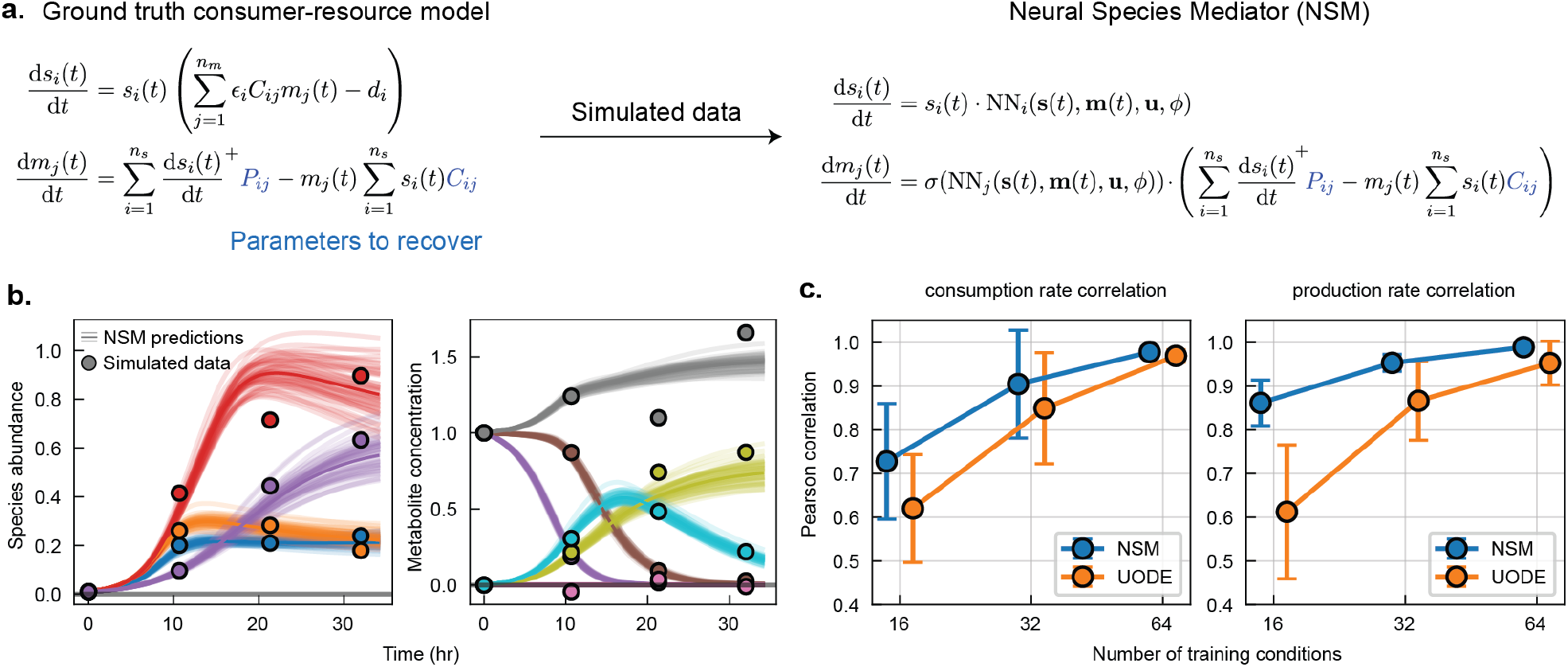
NSM recovers ground-truth consumption and production rates. a. Data of species and metabolites were generated from a ground-truth consumer resource model. Ground truth simulated data was used to fit the NSM, where consumption and production rates were estimated. b. An example of the NSM’s fit to the simulated data, where data points are shown as circles and NSM model predictions are solid lines. Each model prediction represents a draw from the posterior predictive distribution. c. Evaluation of the Pearson correlation between the estimated consumption and production rates and the ground truth values. The amount of training data varied from 16, 32, and 64 different initial conditions, each with 3 measurements taken over a course of 32 hours. Training data was randomly sampled 10 times. The plot shows the mean and standard deviation in the Pearson correlation over the 10 random trials.

To further investigate NSM’s ability to identify direct interactions between species and metabolites, we fit the NSM to combined data set from Clark et al. 2021 [12] and Baranwal et al. 2022 [3] (Fig. 6a). Among the 25 species, 5 species harbor the butyrate production pathway, *Anaerostipes caccae* (AC), *Roseburia intestinalis* (RI), *Coprococcus copri* CC, *Eubacterium rectale* ER, and *Faecalibacterium prausnitzii* (FP) [12, 30]. Consistent with this prior knowledge about the system, the NSM confidently identifies positive butyrate production coefficients to AC, RI, CC, and ER and moderate evidence of butyrate production by FP and *Collinsella aerofaciens* (CA) (Fig. 6b). Butyrate production coefficients identified by the UODE were less constrained with positive values assigned to AC, ER, and *Eggerthella lenta* (EL), which is not a butyrate producer (Fig. S4). While EL does not produce butyrate directly, it displayed a positive impact on butyrate production potentially due to the pH buffering activity of the arginine dihydrolase pathway [28]. Additionally, the UODE did not identify any lactate or succinate producers, while the NSM identified a known lactate producer, *Bifidobacterium pseudocatenulatum* (BP) [18], and several known succinate producers including *Bacteroides vulgatus* (BV) [16], *Bacteroides thetaiotamicron*(BT) [16], and *Prevotella copri* (PC) [22]. These results demonstrate how the multiplication of a neural network with a mechanistic component in the NSM improves the relevance of mechanistic parameters compared to the additive structure of the UODE (Fig. S4).

**Figure 6:**
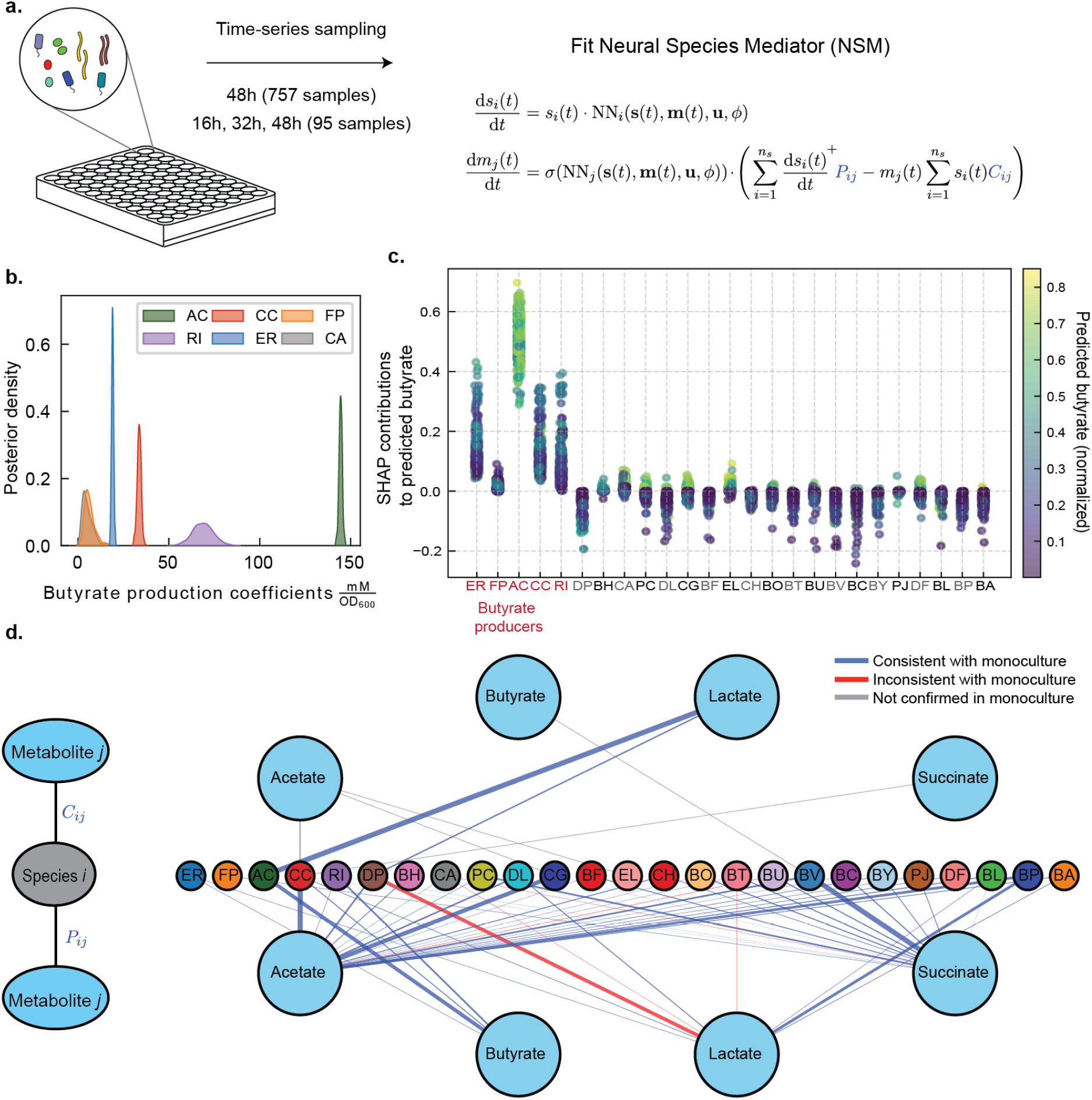
Interpretability of NSM model parameters. a. Experimental data of species abundances and concentrations of acetate, butyrate, lactate, and succinate from coculturing combinations of 25 health-relevant human gut bacteria was used to train the NSM. Estimated consumption and production parameters provide insights into the rate that each species consumes or produces metabolites. b. The posterior parameter distribution of butyrate production coefficients in the NSM show confidently identified butyrate production coefficients of known butyrate producers. The distributions of the six parameters with the largest means are shown, with AC having the largest mean production co-efficient. c. Categorial scatter plot of SHAP values explaining the contribution of species to the NSM’s predicted butyrate concentration. Each column represents the SHAP contributions of each species as predicted over all conditions in the training data set, where the color represents the total predicted butyrate in each condition. d. Interaction network identified by the NSM where connections from metabolites to species represent consumption rate parameters, *C*_*ij*_, and connections from species to metabolites represent production rate parameters, *P*_*ij*_. Edges are colored blue if the trend agrees with empirical evidence from monoculture data, red if the trend was not consistent with monoculture data, and grey if the interaction is not possible to evaluate from monoculture data.

To compare insights from the mechanistic component of NSM to explainable artificial intelligence approaches, we use SHAP to compute the contributions of each species to the NSM’s predictions of butyrate across all training conditions (Fig. 6c). SHAP identified positive contributions of several species that are not butyrate produces such as CA, EL, and *Prevotella copri* (PC). While EL does not produce butyrate directly, it displayed a positive impact on butyrate production potentially due to the pH buffering activity of the arginine dihydrolase pathway [28]. Both CA and EL were significantly positively correlated with pH in Clark et al. 2021 [12]. While SHAP values cannot be interpreted as directly causal interactions, this method can uncover species that contribute both directly and indirectly to predicted functions. Therefore, these interpretation approaches are complementary since the discovery of influential indirect interactions is not possible to identify from the analysis of the mechanistic model parameters.

The full set of mechanistic model parameters in the NSM can be used to generate an interaction network based on species consumption and production rates of metabolites (Fig. 6c). To corroborate these results, the metabolite production and consumption profiles of monocultures can provide insight into direct interactions between species and metabolites. Interactions that are consistent between monoculture data (i.e. significant increases or decreases in metabolite concentration from inoculation to endpoint based on a one sample one sided t-test) and NSM are shown as blue edges, edges that are inconsistent with monoculture are red, and edges that cannot be confirmed are grey (e.g., consumption of acetate, butyrate, and succinate cannot be determined since the media was not supplemented with these metabolites). Inspection of this network reveals a confirmatory interaction where AC consumes lactate and produces butyrate [12].

However, the model identifies that *D. piger* (DP) produces lactate, which is a false positive as numerous studies have shown DP consumes lactate rather than produces this metabolite [34, 42]. Inspection of the endpoint abundances of AC in conditions that were co-cultured with DP reveal a significant decrease in abundance compared to conditions that did not include DP (t-test *p <* 1 *×* 10^*−*5^, Fig. S6a). The concentration of lactate at 48 hours was also significantly higher in conditions with both AC and DP compared to conditions with only AC, suggesting that DP inhibits AC’s ability to consume lactate (t-test *p <* 1 *×* 10^*−*5^, Fig. S6b). Further, the distribution of DP’s SHAP contribution to AC was negative in more than 99% of conditions where both DP and AC were co-cultured (Fig. S6c). We therefore hypothesize that DP inhibits the growth of AC on lactate, resulting in an increase in lactate concentration in communities that include both DP and AC. Consequently, the model incorrectly estimates that DP directly produces lactate. Misidentified parameters may be due to limited training data based on our analysis of the NSM’s accurate recovery of ground truth parameters that improved as training data size increases (Fig. 5c). In sum, interpreting mechanistic model parameters is complementary to explainable artificial intelligence methods such as SHAP ([45]). These methods can be combined to reveal direct causal relationships and other indirect biologically relevant interactions.

## Discussion

To unlock the potential of microbial communities, it is essential to develop predictive models that not only capture their dynamic behavior and emergent functions but also integrate physical constraints and provide mechanistic insights. Predicting the dynamics of species and metabolites in synthetic microbial communities is a challenge, as mechanistic models often fail to capture system complexity and machine learning methods can make physically unrealistic predictions. Further, existing machine learning frameworks that have been applied to synthetic microbial communities are typically based on recurrent neural network architectures [3, 51], which make predictions at fixed time intervals. In this study, we develop and validate a continuous time machine learning model based on a neural ordinary differential equation architecture that incorporates a mechanistic model of metabolite dynamics to enforce physical constraints. The resulting NSM model outperforms both mechanistic models and machine learning models in prediction of metabolite concentrations, especially when the amount of training data is limited (Fig. 4).

The augmentation of mechanistic models with neural networks has been applied in a variety of disciplines [24, 40]. While previous studies have incorporated neural networks by adding a neural network to a mechanistic model [1], we show that the multiplicative structure preserves physical constraints, improves model prediction performance and the accuracy of estimates of mechanistic model parameters. While the proposed model performs well on several experimental datasets, the mechanistic component may need to be modified depending on the system. For example, we assume that the rate of metabolite production is proportional to species growth rates. However, this assumption can be modified in cases where species release metabolites in stationary phase [23, 48]. In other applications, the multiplication of a strictly positive neural network with a mechanistic model could be a general framework to produce a flexible class of physically constrained ordinary differential equation models.

As a continuous-time model, the NSM can train on data with irregular sampling intervals (Fig. 2, 4) and can make predictions over a continuous time range. While this enables capabilities for interpolation and extrapolation that are not possible using RNNs, ordinary differential equation models require additional computational costs compared to discrete-time models because of numerical integration. Despite this limitation, parameter estimation including uncertainty quantification of the NSM took less than eight hours on a relatively large synthetic microbial community data set with 757 initial combinations of 25 species (over 13,000 measurements of species and metabolites) [12]. Further, it is possible to misidentify interactions between species and metabolites in cases where species indirectly impact metabolite dynamics via interactions with other species. In particular, the NSM identified DP as a lactate producer due to the elevated levels of lactate in conditions with both DP and AC (Fig 6c). Therefore, the inferred parameters should be used as a hypothesis generating tool for further investigation.

To provide confidence in identified interactions, we use Bayesian parameter inference to quantify uncertainty in interaction parameters. We used a computationally scalable variational inference approach to estimate the posterior parameter distribution as an independent Gaussian for each parameter. Future work could extend this approach using a more flexible variational posterior such as a multivariate Gaussian to capture correlations and improve uncertainty estimates. An additional area of future work could be to analyze the activity of the neural network to gain insight into cases where the mechanistic component is insufficient. Neural network architectures such as Kolmogorov-Arnold networks (KANs) [29] may be a promising alternative that provides flexibility and greater interpretability compared to the feed-forward neural network used in this study. The mechanistic component of the model could also be further refined to force additional constraints such as conservation of mass and energy [35]. Finally, microbial communities may be influenced by unmeasured and unknown metabolites. In this case, an avenue of future research is augmenting NSM with dynamic latent variables for additional flexibility [14]. The latent variables could represent additional metabolites in the model with initial conditions that become additional model parameters.

Embedding a mechanistic model into a neural ordinary differential equation produces a physically constrained model that can capture state-dependent shifts in metabolite production and consumption in microbial communities. We used a Bayesian inference approach to enable uncertainty quantification in model parameters and showed how the incorporation of physical constraints increases prediction accuracy and reduces parameter uncertainty. The ability to quantify model uncertainty can be leveraged for Bayesian experimental design, where experimental conditions are selected based on the expected information gain [51]. We envision that this model will play a critical role in accelerating the design-test-learn framework for the discovery of optimized synthetic microbial communities for a broad range of applications spanning precision medicine, sustainable agriculture, environmental cleanup and biomanufacturing and their governing molecular and ecological principles [27].

## Methods

### The consumer-resource model with growth-associated metabolite production

To generate the simulated data used in Fig. 5, Fig. S1, and Fig. S2, we used a consumer-resource model with growth-associated metabolite production,

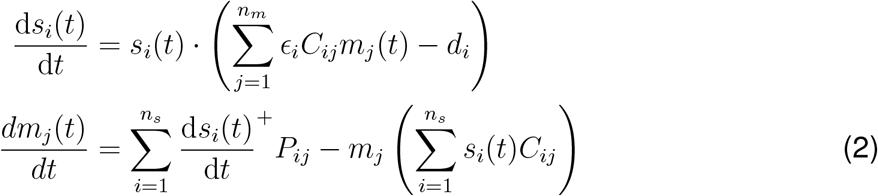

where *ϵ*_*i*_ is the efficiency that species *i* grows on resources, *d*_*i*_ is the death rate of species *i, C*_*ij*_ is the consumption rate of species *i* on metabolite *j*, and *P*_*ij*_ is the production rate of metabolite *j* by species *i*. Species growth efficiencies were randomly sampled as *ϵ*_*i*_ *∼ 𝒰* [.8, .9] where *𝒰* represents the uniform distribution. Death rates were randomly sampled as *d*_*i*_ *∼ 𝒰* [.008, .012]. Ground truth production rates where randomly generated as *P*_*ij*_ *∼ 𝒰* [0, 2] *·* Bernoulli(.5), where Bernoulli(.5) is a random variable with equal probability of being either zero or one. Each consumption rate was initially sampled as *C*_*ij*_ *∼ 𝒰* [.5, 1]. Consumption rates were then set to zero in cases that caused positive feedback loops in metabolite production and consumption. Simulated data was generated by randomly sampling combinations of eight species and six resources where initial species abundances were set to .01 and initial resource concentrations were 1. The data included species abundances and resource concentrations sampled at 10.6, 21.3, and 32 hours, where data was corrupted with zero-mean Gaussian noise with variance *σ*^2^(*x*) = .001 + .01 *· x*^2^, where *x* represents simulated species or metabolite.

### The ecological neural ordinary differential equation (eNODE) and neural species mediator (NSM) models

#### Neural species mediator architecture

We denote the time-dependent vector of species abundances as 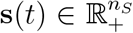, metabolites as 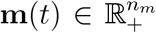, and additional system inputs as 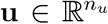. The neural species mediator (NSM) is given by the following set of coupled differential equations,

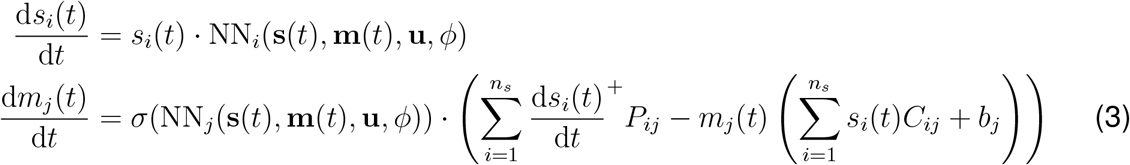

where *P*_*ij*_ *∈* ℝ_+_ is the strictly-positive growth-associated rate that species *i* produces metabolite *j, C*_*ij*_ *∈* ℝ_+_ is the strictly-positive rate that species *i* consumes metabolite *j*, and *b*_*j*_ *∈* ℝ_+_ is the constant degradation rate of metabolite *j*. We use the rectified linear unit function, *a*^+^ = max(0, *a*), to model metabolite production as proportional to species growth. The structure of the mechanistic component can be modified as follows to account for biomass associated metabolite production where metabolites can be produced during stationary and death phase,

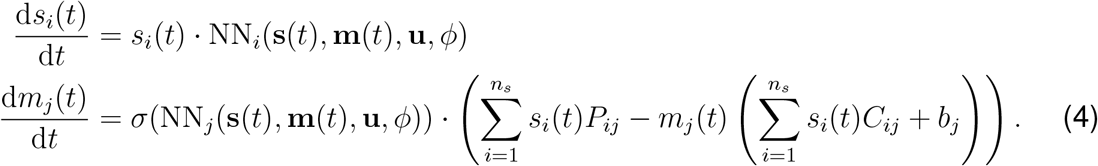

#### Neural network architecture

The neural network in the eNODE and NSM (Eq. 1) uses a feedforward architecture to predict the rate of change of species and metabolites,

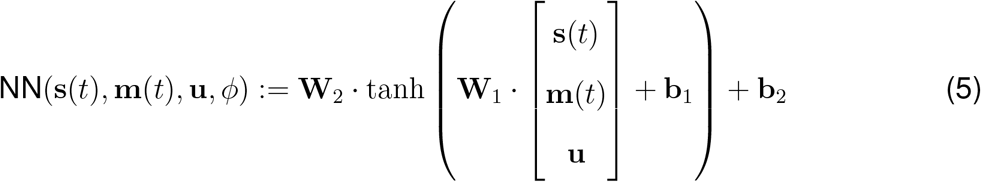

The set of weights and biases in the neural network is denoted as *ϕ* = {**W**_1_, **b**_1_, **W**_2_, **b**_2_}.

#### Constraining mechanistic model parameters

The parameters in the consumer-resource model are all strictly non-negative and represent rates of metabolite production, consumption, and degradation. To ensure non-negativity of rate parameters, we apply the SOFTPLUS function with base ten, *σ*_10_(*a*) = log_10_(1 + 10^*a*^), to each parameter. We denote the unconstrained parameter using a hat, e.g. *C*_*ij*_ = *σ*_10_(*Ĉ*_*ij*_). The full set of parameters in the NSM is denoted as 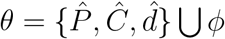.

### Parameter estimation

#### General description of data structure and model predictions

The central modeling goal in this study is to predict the distribution of experimental dependent variables given all independent variables. In this case, dependent variables include species abundances and metabolite concentrations and independent variables include initial conditions of species and metabolites, media conditions such as the presence or absence of resources, and the measurement time. We denote an experimental condition as the set of independent variables, *x* = {**s**(0), **m**(0), **u**, *τ*}. We denote the vector of measured dependent variables as

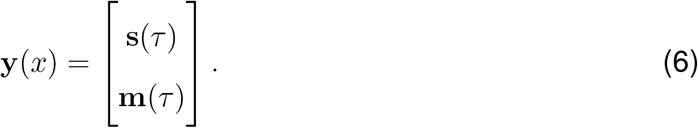

We denote model predictions of the dependent variables as **ŷ**(*x*),

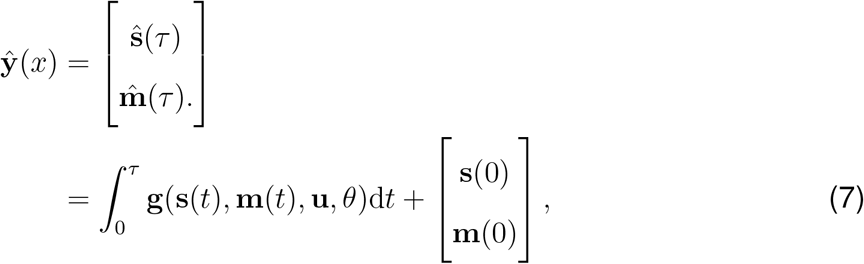

where **g**(**s**(*t*), **m**(*t*), **u**, *θ*) is a model such as the NSM, eNODE, gLV, or CR, that predicts rate of change of species and metabolites. A data set is composed of measured species abundances and metabolite concentrations in *n* unique experimental conditions, which we denote as *𝒟* = {**y**(*x*_1_), …, **y**(*x*_*n*_)}.

#### Maximum likelihood

Prediction of species abundances and metabolite concentrations is a regression problem that involves optimizing model parameters such as species growth rates, metabolite consumption rates, and neural network weights and biases, such that the error between model predictions and measured values is minimized. Experimental measurements of species and metabolites are inherently noisy, meaning that repeated measurements of the same variable under the same experimental conditions will result in a distribution of values. In practice, this distribution is approximately Gaussian with a standard deviation that scales linearly with the magnitude of the measured variable. We therefore model the conditional distribution of an experimental measurement in a particular experimental condition as a Gaussian with a mean and variance predicted by the model,

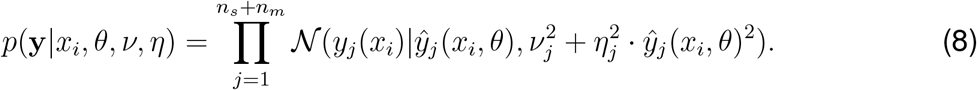

Given this model for the distribution of a measured variable, the likelihood of a set of independent measurements is

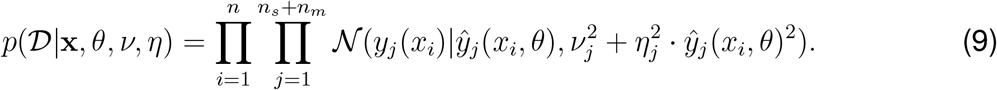

Maximizing the likelihood of the data with respect to parameters is equivalent to maximizing the log likelihood and gives the maximum likelihood estimate (MLE),

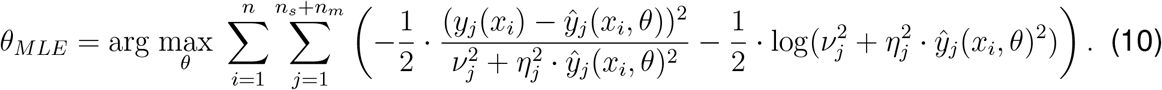

In the case where measurement noise is assumed constant, the maximum likelihood estimate is equivalent to minimizing the sum of squared error between model predictions and measured values,

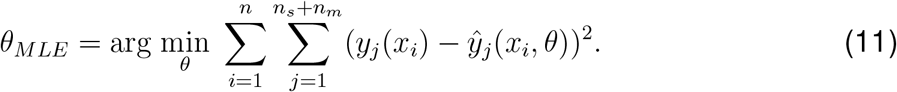

#### Maximum A Posteriori

Maximum likelihood parameter estimation can suffer poor generalization due to the model fitting to measurement noise. To mitigate this, a prior on parameters centered at zero promotes simplicity in the model. Furthermore, using an independent Gaussian distribution centered at zero for each parameter prior promotes sparsity by driving the posterior distribution of unnecessary parameters to the prior mean [4]. We use a Gaussian prior centered at zero with a variance for each parameter,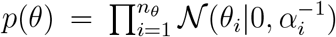. Using Bayes’ theorem gives the posterior probability of parameters,

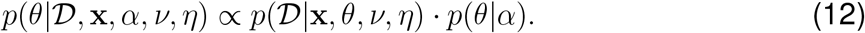

Maximizing the posterior probability gives the maximum a posteriori (MAP) estimate,

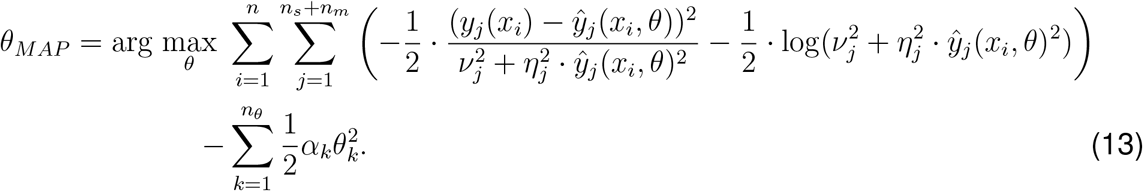

#### Variational inference

The goal of Bayesian inference is to determine the conditional probability distribution of model parameters given a data set, which is termed the posterior parameter distribution. Variational inference is a method that casts estimating the posterior parameter distribution as an optimization problem where the objective is to minimize the Kullback-Leibler divergence between an approximate parametric probability density function and the true parameter posterior. A popular choice for a probability density function is an independent Gaussian for each parameter due to its analytical tractability and computational scalability, denoted here as 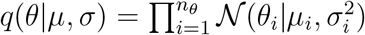. To enable a more compact notation we denote the posterior as *p*(*θ*|*𝒟, α, ν, η*), omitting the dependence on the set of experimental conditions, **x**. The Kullback-Leibler divergence between the approximate posterior and the true posterior is

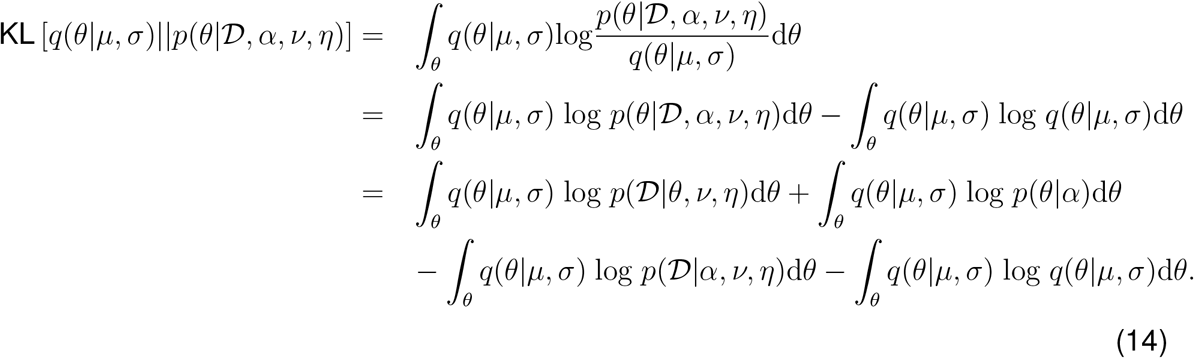

Keeping only the terms that depend on the variational parameters *µ* and *σ*, we define

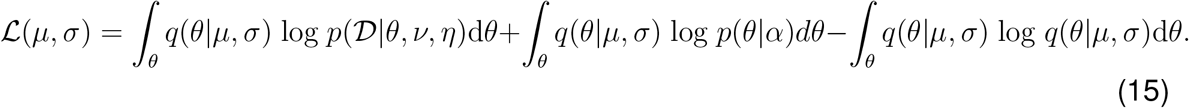

The function *ℒ* (*µ, σ*) is called the evidence lower bound (ELBO). Although the expectation of the log likelihood is analytically intractable, the ELBO can be estimated using a Monte Carlo approximation,

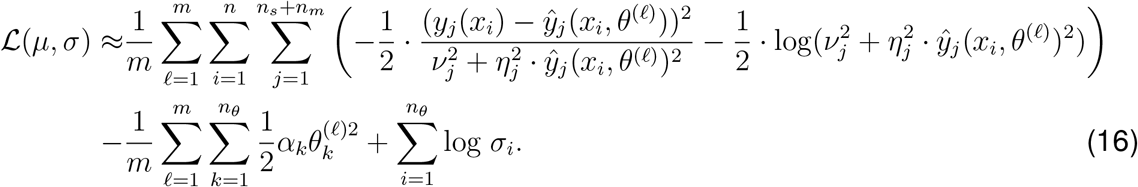

where *θ*^(*ℓ*)^ *∼ q*(*θ*|*µ, σ*). Optimization of the ELBO with respect to *µ* and *σ* is performed using the ADAM algorithm with the default parameters of *α* = .001, *β*_1_ = .9, and *β*_2_ = .999 [25]. The slope of a linear fit to the ELBO over the previous 100 training epochs is used to evaluate convergence. The algorithm terminates when the slope of the ELBO is less than .001 for five consecutive evaluations.

### Expectation Maximization to optimize hyperparameters

Model hyperparameters include the precision of the parameter prior, *α*, and the measurement noise variance parameters *ν* and *η*. The expectation maximization (EM) algorithm iterates between inference of the parameter posterior distribution and then maximization of the expected joint distribution of the data and model parameters with respect to model hyperparameters [4]. Because inference of the posterior depends on the hyperparameters, the posterior distribution needs to be updated given the optimized hyperparameters. The process of iterating between updating the posterior and updating hyperparameters continues until convergence of the ELBO. The expectation of the joint distribution of the data and model parameters is

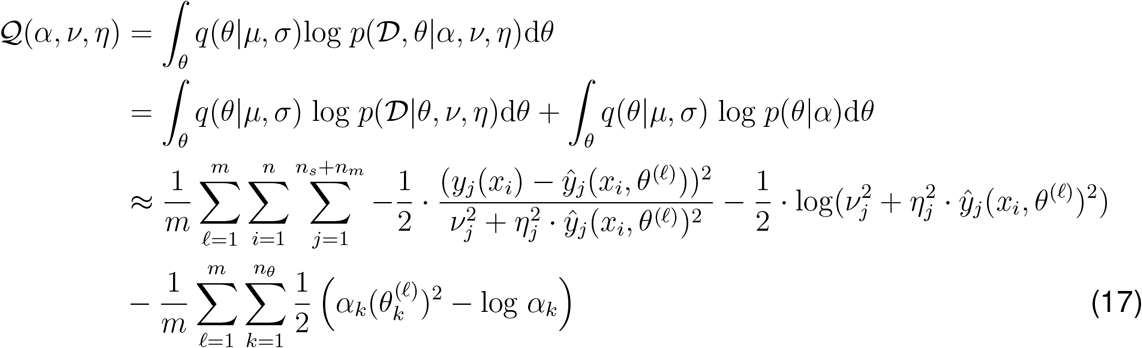

where *θ*^*j*^ *∼ 𝒩* (*θ*|*µ, σ*). Maximization of *𝒬* (*α, ν, η*) with respect to *α* gives the update equation,

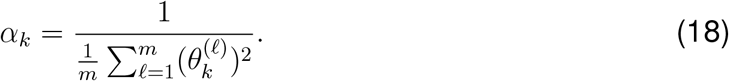

*𝒬* (*α, ν, η*) is maximized with respect to *ν* and *η* when

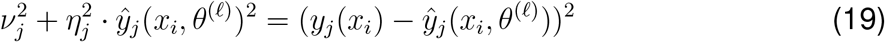

for each *j* = 1, …, *n*_*s*_ + *n*_*m*_ for all *ℓ* = 1, …, *m* and *i* = 1, …, *n*.

### Models for comparison with the NSM

#### Generalized Lotka-Volterra (gLV)

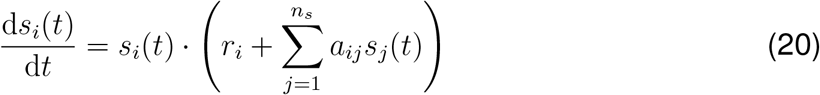

#### Consumer-resource (CR)

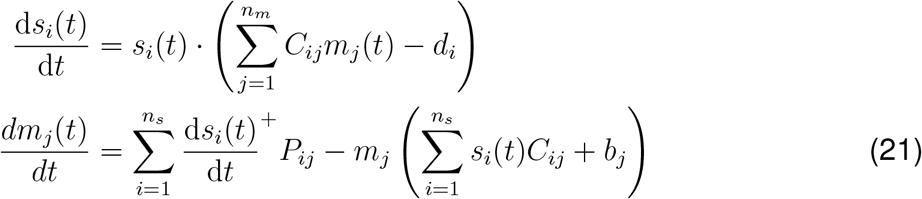

#### Universal ordinary differential equation (UODE)

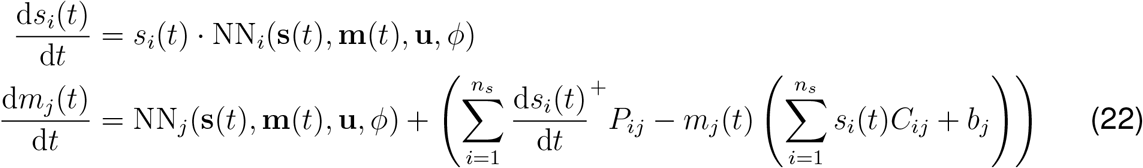

### Strain culturing and hydrogen sulfide quantification

Strain maintenance and culturing of *Faecalibacteirum prausnitzii* (FP) DSMZ 17677 and *Desulfovibrio piger* (DP) ATCC 29098 were performed according to Clark 2021 [12]. Side-by-side monoculture and coculture growth kinetics were measured in response to varying key nutrients (glucose, lactate, and sulfate) according to 3-factor 3-level full factorial experimental design. The variable nutrients were assembled according to the experimental design using a Tecan Evo Liquid handling robot (see Supplemental data file S?? for base medium composition and variable component design levels). The media used were based on that developed in Clark 2021 [12]. Replicate plates copies corresponding to each sample timepoint were prepared, inoculated, and cultured in parallel, with one plate harvested at each timepoint. This was done so as to avoid depletion of culture volume across multiple measurements from small volume cultures and to avoid disturbing apparent three-dimensional structure of the cultures (which we expected may be contribute to interspecies-interactions). Identically prepared cultures were each harvested at 7, 10, 19, and 37 hour timepoints, each containing n=3 biological replicates of each media-species or media-coculture combination. Measurements of all biological replicates are included in Supplementary Data 1. Inoculation density for monoculture conditions was .01 OD600 and coculture experiments was .005 OD600 per species (.01 OD600 total). Hydrogen sulfide production was measured at each timepoint using a 96-well plate-based adaption of the cline assay according to Hromada 2023 [21]. Coculture relative abundance was measured by 16S rRNA sequencing on an illumine MiSeq according to detailed methods in Clark 2021 [12]. Total growth was measured by optical density at 600nm, diluting and back calculating if greater than the linear range of the instrument (1.0 OD600).

### Correlated coefficients z-test

We use the correlated coefficients z-test to test whether there is a significant difference in correlations between measured and predicted values of two competing models [36]. The test yields a z-score testing the null hypothesis of equal correlation coefficients, accounting for correlations between the two model predictions.

## Data and code availability

All code and data used for model training and validation will be publicly available at https://github.com/VenturelliLab/Thompson_et_al_2025 upon acceptance of the manuscript for publication. Code is provided to the reviewers as a Supplementary File.

## Author contributions

J.T., V.M.Z, and O.S.V. conceived the study. B.M.C. carried out experiments. J.T. performed computational modeling. J.T. and B.M.C. performed statistical analyses of experimental data. J.T., V.M.Z., and O.S.V. analyzed data. J.T. and O.S.V. wrote the paper and all authors provided feedback on the paper. O.S.V. secured funding.

## Acknowledgments

This work was supported by the Defense Advanced Research Projects Agency (DARPA) under Contract No. HR0011-23-2-0001, HR0011-25-3-0072 and the National Institutes of Health R01EB030340 to Ophelia S. Venturelli. Victor M. Zavala acknowledges support from the National Science Foundation under award CBET-2315963.

