## Supplementary Information for "Physics-constrained neural ordinary differential equation models to discover and predict microbial community dynamics"

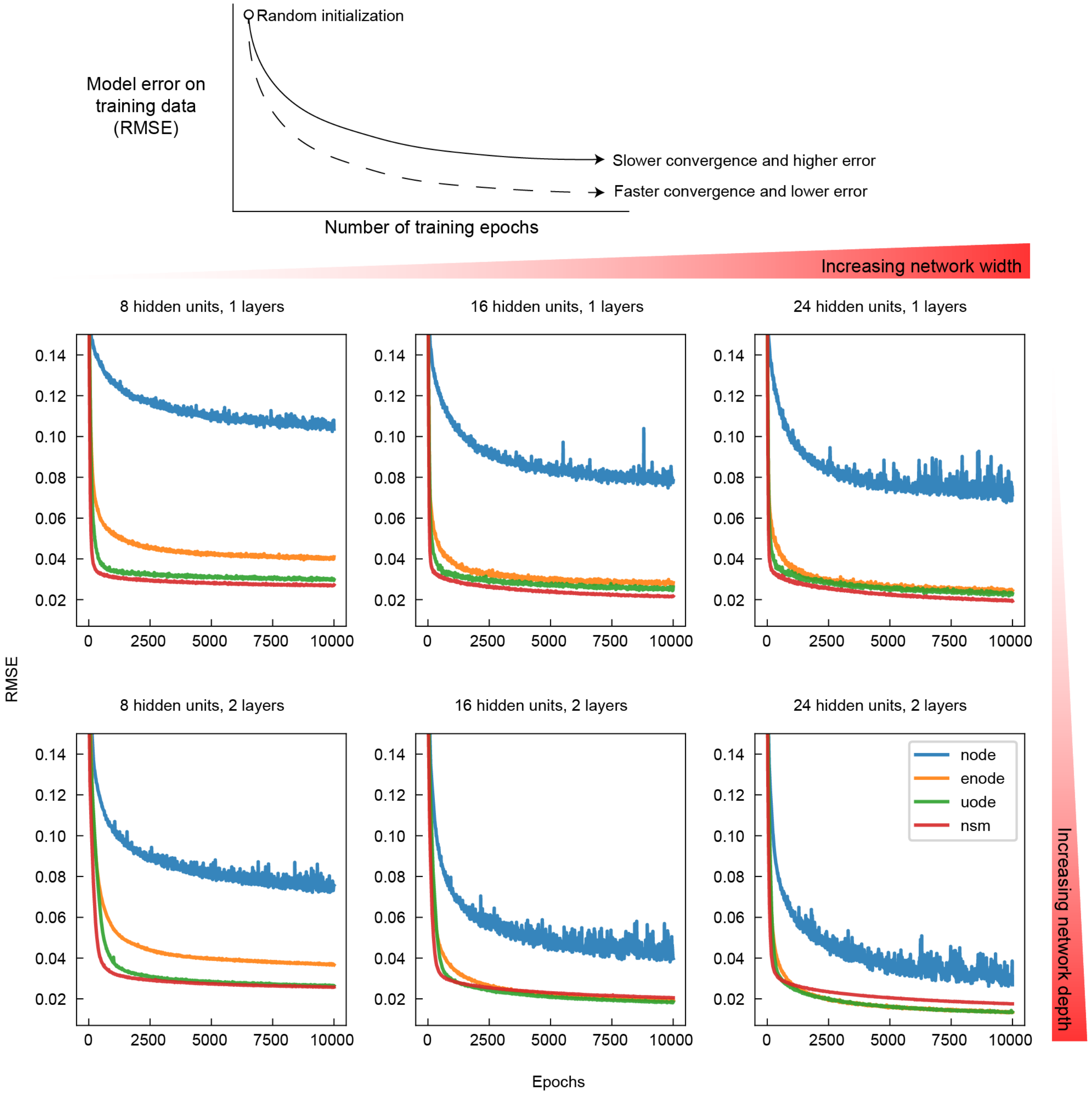


Figure 1: **Comparison of training progress on simulated data** Training progress on the FP-DP-Sulfide data set was evaluated using the root-mean-squared-error (RMSE) and compared between the NODE, eNODE, UODE, and NSM models. Simple neural network architectures result in poor convergence by the NODE due to the inability to predict species absence for species that were not inoculated. Similarly, the eNODE requires a more flexible neural network architecture to learn metabolite dynamics. Increasing the complexity of the neural network by increasing the number of hidden layers and hidden units is required for both the NODE and eNODE to match the fit of the UODE and NSM.

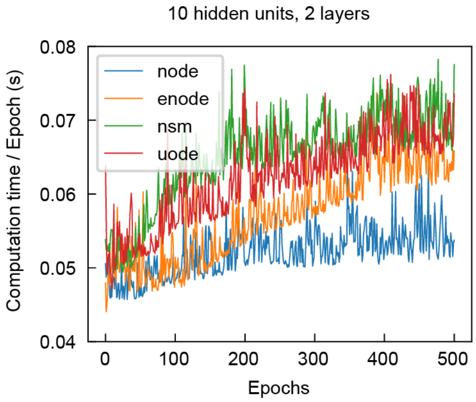


Figure 2: **Comparison of computation time per epoch.** The NSM and UODE models take slightly longer to train per epoch compared to eNODE and NODE and generally training time per epoch increases as training proceeds.

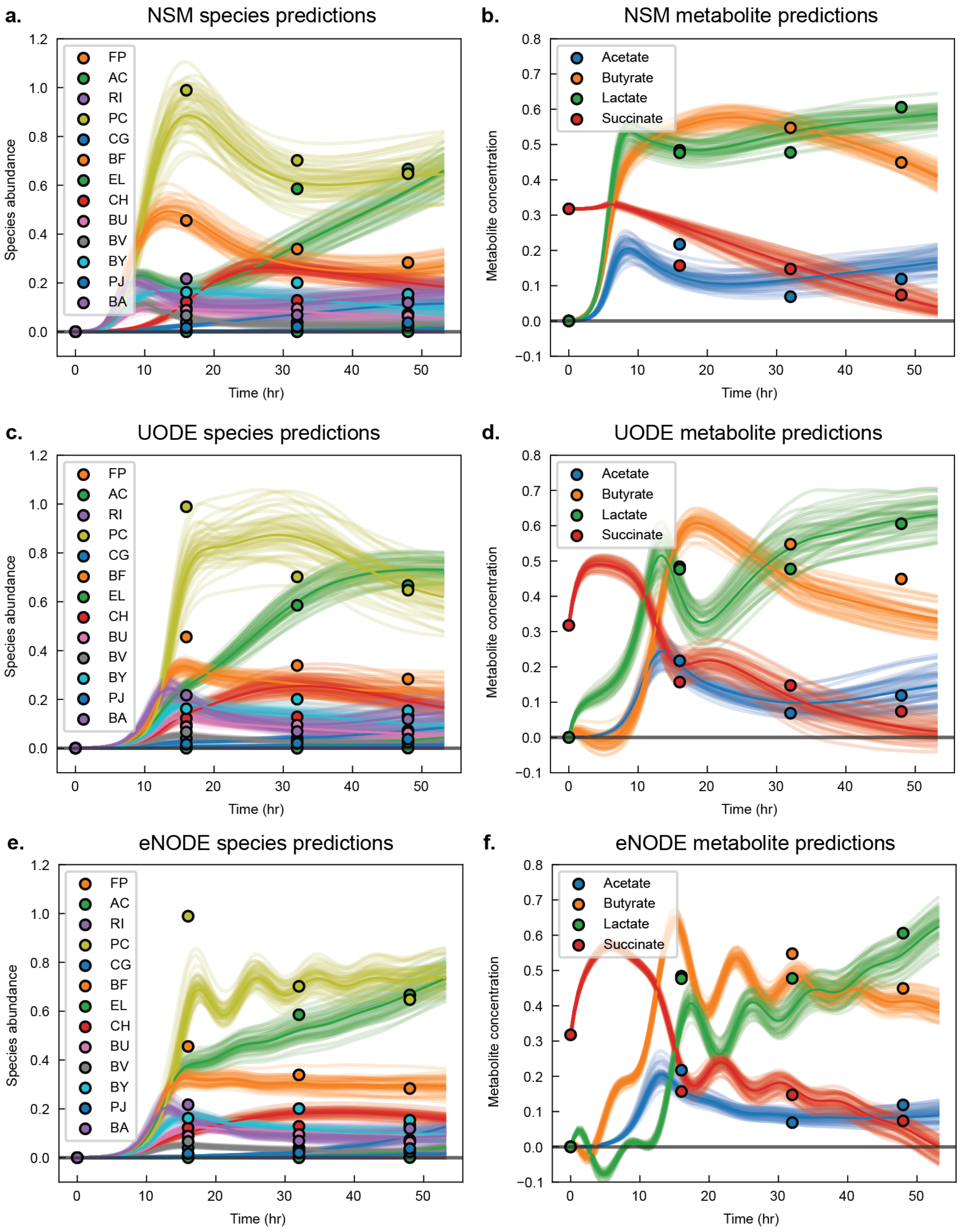


Figure 3: **Comparison of NSM, UODE, and eNODE species and metabolite predictions**

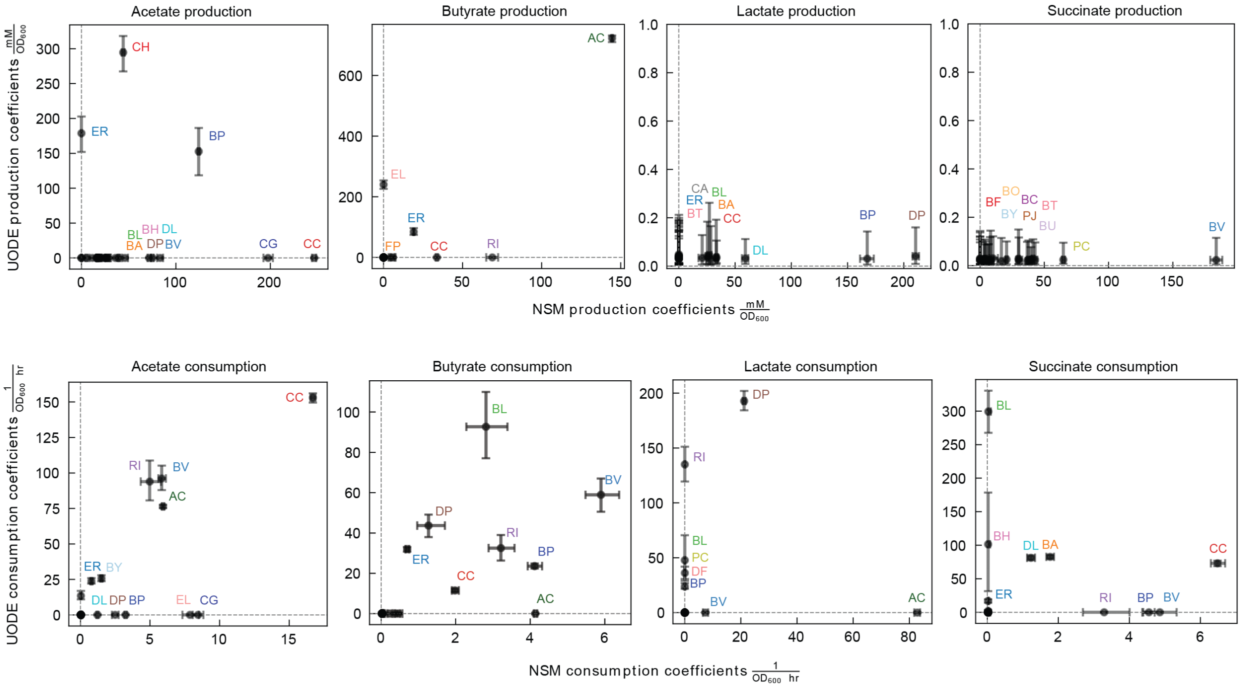


Figure 4: **Posterior parameter distribution of consumption and production coefficients in the NSM and UODE models.** Production and consumption parameters of the mechanistic component of the UODE model are plotted versus the NSM model parameters. Each point shows the median and interquartile range of the posterior parameter distribution. Production rates for each metabolite are plotted in the top row and consumption rates are plotted in the bottom row.

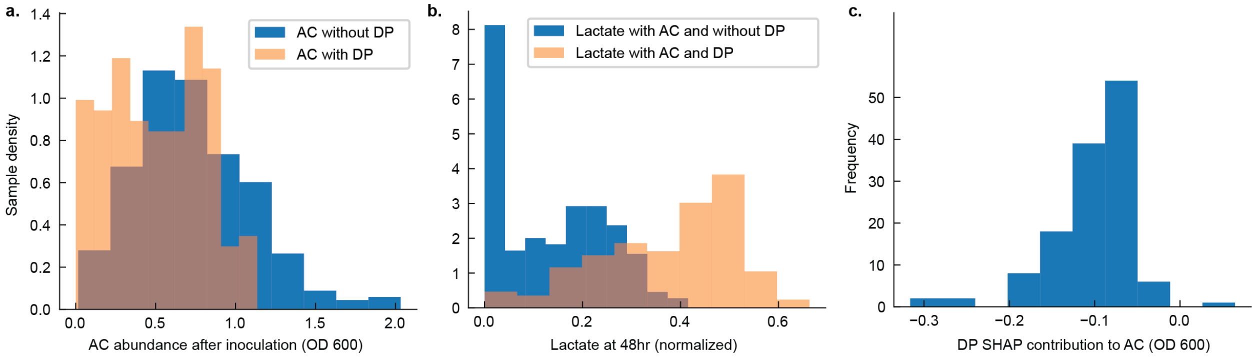


Figure 5: **Inhibition interaction of DP on AC** a. The abundance of AC (OD 600) in conditions without DP is significantly greater than the abundance of AC in conditions when co-cultured with DP (t-test, p-value $< 1 \times 10^{-5}$). b. The concentration of lactate at 48 hours in conditions with AC and without DP is significantly lower than the concentration in conditions when AC is co-cultured with DP (t-test, p-value $< 1 \times 10^{-5}$). c. The SHAP contribution of DP on NSM predictions of AC in conditions that contain both AC and DP is negative in more than 99% of conditions.
